# Single kinetochores execute an ordered series of molecular events as the Spindle Assembly Checkpoint is silenced

**DOI:** 10.1101/2025.05.23.655733

**Authors:** Catriona C. Conway, Tsvetelina Germanova, Sara Toral-Perez, Catherine Coates, Jonathon Pines, Nigel J. Burroughs, Andrew D. McAinsh

## Abstract

The Spindle Assembly Checkpoint (SAC) delays anaphase onset until all kinetochores are stably attached to microtubules, thus promoting error-free chromosome segregation. Multiple molecular events are implicated in SAC silencing, including removal of phospho-marks, protein (un)binding, and structural reorganisation of the kinetochore – but we currently lack a quantitative map of how these events unfold through time. Here, we use the levels of the checkpoint protein MAD2 to create a pseudo-timeline of SAC silencing at single kinetochores. We demonstrate how silencing proceeds through an ordered series of molecular events where MAD2-Spindly unbinds first and then the KNL1 catalytic platform disassembles, with a pool of active MPS1 retained. Coincidently, the NDC80 ensemble reconfigures in response to high microtubule occupancy. Kinetochores next switch into a mature attachment state that then undergoes gradual further stabilisation through NDC80 tail dephosphorylation. By preventing biorientation, we also define otherwise hidden kinetochore states involved in error correction cycles. This includes a “poised” state which we propose allows for error correction and rapid reactivation of the SAC. These results provide a critical temporal framework for understanding the mechanisms of SAC silencing and error correction at single human kinetochores.

## Introduction

The spindle assembly checkpoint (SAC) is a dynamic surveillance mechanism that operates to limit chromosome mis-segregation during cell division (McAinsh and Kops, 2023). This is important because changes in chromosome number, even by one, can lead to aneuploidy, which is associated with multiple human pathologies and developmental disorders (Chunduri and Storchová, 2019; Santaguida and Amon, 2015; Sdeor et al., 2024). The SAC monitors the attachment of each chromosome to the microtubule-based mitotic spindle. Such attachments are mediated by kinetochores, multi-protein complexes that assemble on the centromere of each chromosome (Musacchio and Desai, 2017). If a kinetochore is not bound to microtubules, it generates a diffusible signal (MCC, mitotic checkpoint complex) which can inhibit the Anaphase promoting complex/cyclosome (APC/C) – this is a SAC “ON” state (McAinsh and Kops, 2023). In this way, even a single unattached kinetochore can be sufficient to prevent the onset of anaphase and limit potentially inaccurate chromosome segregation.

In human cells, all kinetochores begin life unattached as the nuclear envelope breaks down, and thus the default state of the SAC is “ON”. How the MCC is generated at kinetochores has been intensely investigated and in metazoans involves 1) activation of MPS1 kinase (Kuijt et al., 2020), 2) phospho-dependent recruitment of BUB3, BUB1, MAD1 and CDC20 to core kinetochore scaffolds (Fischer et al., 2022; Ji et al., 2017; London et al., 2012; Primorac et al., 2013; Shepperd et al., 2012; Yamagishi et al., 2012; Zhang et al., 2017), 3) the assembly of multiprotein complexes that function as a catalytic platform which captures soluble MAD2 and releases a complex of MAD2-BUBR1-BUB3-CDC20 (the MCC; Faesen et al., 2017; Ji et al., 2017; London and Biggins, 2014; Piano et al., 2021), and 4) RZZ/Spindly recruitment of MAD1-MAD2 to the corona, potentially forming a second binding site for soluble MAD2 independent of KNL1 (Allan et al., 2020; Buffin et al., 2005; Gassmann et al., 2010; Kops et al., 2005; Rodriguez-Rodriguez et al., 2018; Silió et al., 2015). As microtubule-kinetochore interactions occur, MCC production must be extinguished and the above events reversed (moving to a SAC “OFF” state). This involves dephosphorylation of kinetochore proteins, driven by a shift in kinase-phosphatase feedback loops (Saurin, 2018), and the physical stripping of MAD1-MAD2 from kinetochores by Dynein molecular motors (Famulski et al., 2011; Gassmann et al., 2010; Howell et al., 2001; Ide et al., 2023; Silva et al., 2014). SAC silencing also correlates with mechanical events, including changes in intra-kinetochore tension and conformation (Maresca and Salmon, 2009; Renda et al., 2020; Roscioli et al., 2020; Uchida et al., 2009; Uchida et al., 2021) and inter-sister tension (Itoh et al., 2018; Maresca and Salmon, 2009; Uchida et al., 2009; Yoo et al., 2018). What remains unclear is whether kinetochores simply switch from a SAC “ON” to a SAC “OFF” state, or whether there are intermediate steps that reflect different functional states of the kinetochore, i.e. is there a defined sequence of events that proceed in order.

The SAC silencing mechanism faces two complications. Firstly, human kinetochores assemble from multiple protein copies to form an ensemble, which binds multiple microtubules (∼10; Kiewisz et al., 2022; O’Toole et al., 2020). How SAC silencing operates across ensemble kinetochores is a mystery, particularly since several studies suggest that full microtubule occupancy is not a prerequisite for silencing (Dudka et al., 2018; Etemad et al., 2019; Kuhn and Dumont, 2019). Secondly, end-on attachment is required to silence the SAC but not sufficient to ensure accurate chromosome segregation (Etemad et al., 2015; Kapoor et al., 2000; Tauchman et al., 2015). This is because sister kinetochores can form erroneous attachments, e.g. both sisters bound to one pole (syntely) or one sister bound to both poles (merotely). Tension-dependent error correction, mediated by Aurora B, is expected to destabilise erroneous attachments (Carmena et al., 2012; Krenn and Musacchio, 2015; Lampson and Cheeseman, 2011; Lampson and Grishchuk, 2017; McVey et al., 2021), leading to microtubule detachment and reactivation of the SAC. But how error correction and SAC silencing are sequenced at single kinetochores is unresolved.

Here, we develop a quantitative single kinetochore-based fixed-cell approach to probe the molecular events during SAC silencing. This circumvents the current limitations of live-cell imaging (low numbers of kinetochores, limited reporters, especially for phospho-marks, and photobleaching/toxicity). Using kinetochore-bound MAD2 signal to order kinetochores, we reveal that the kinetochore undergoes an ordered series of molecular events as the SAC silences and show that SAC silencing at individual kinetochores precedes the stabilisation of microtubule attachments or activation of error correction. These experiments reveal new functional states of the kinetochore, including a “poised” state which we propose allows for error correction and rapid reactivation of the SAC.

## Results and discussion

### Establishing MAD2 pseudo-time to investigate SAC silencing in human cells

To investigate how molecular events may be sequenced during SAC silencing, we established a method to order single kinetochores based on their levels of bound MAD2, which we used as a proxy for SAC activity (Figure 1A). We hypothesised that kinetochores adapt as SAC silencing proceeds and that kinetochores with different MAD2 levels will reflect different functional states. For this we used human non-transformed RPE1 cells which express Venus-MAD2 from the endogenous locus (Collin et al., 2013). Cells were fixed and stained with anti-CENP-C antibodies to allow automated detection of kinetochore positions and quantification of Venus-MAD2 per kinetochore. As expected, kinetochore-bound MAD2 was highest in early prometaphase and reduced through prometaphase to metaphase, but the amount of MAD2 on individual kinetochores was variable throughout mitosis (Figure 1B). We therefore ranked kinetochores by Venus-MAD2 intensity and assigned them to 25 bins, each representing 4% of all kinetochores, producing a MAD2 pseudo-timeline (black line, Figure 1C). We observed that kinetochores in all mitotic stages were represented at all points in the pseudo-timeline (Figure 1A, lower panel). This single-kinetochore approach therefore avoids mistreating kinetochores in each mitotic stage as being in an equivalent functional state, i.e. a kinetochore can be unattached even though it is in metaphase (Figure 1A, arrow on right panel). To ensure that the pseudo-timeline was robust, we co-stained with antibodies against MAD2 or MAD1-pT716 and assessed their intensities within the MAD2 pseudo-timeline bins created by endogenously labelled Venus-MAD2 (Figure 1D). These curves closely followed each other, with correlation coefficients of Venus-MAD2 versus anti-MAD2 and anti-MAD1-pT716 antibodies of 0.8693 and 0.8684, respectively. To enable comparison with other molecular events, we then defined the “half-change” point as the MAD2 pseudo-time where signal has changed by half.

**Figure 1:**
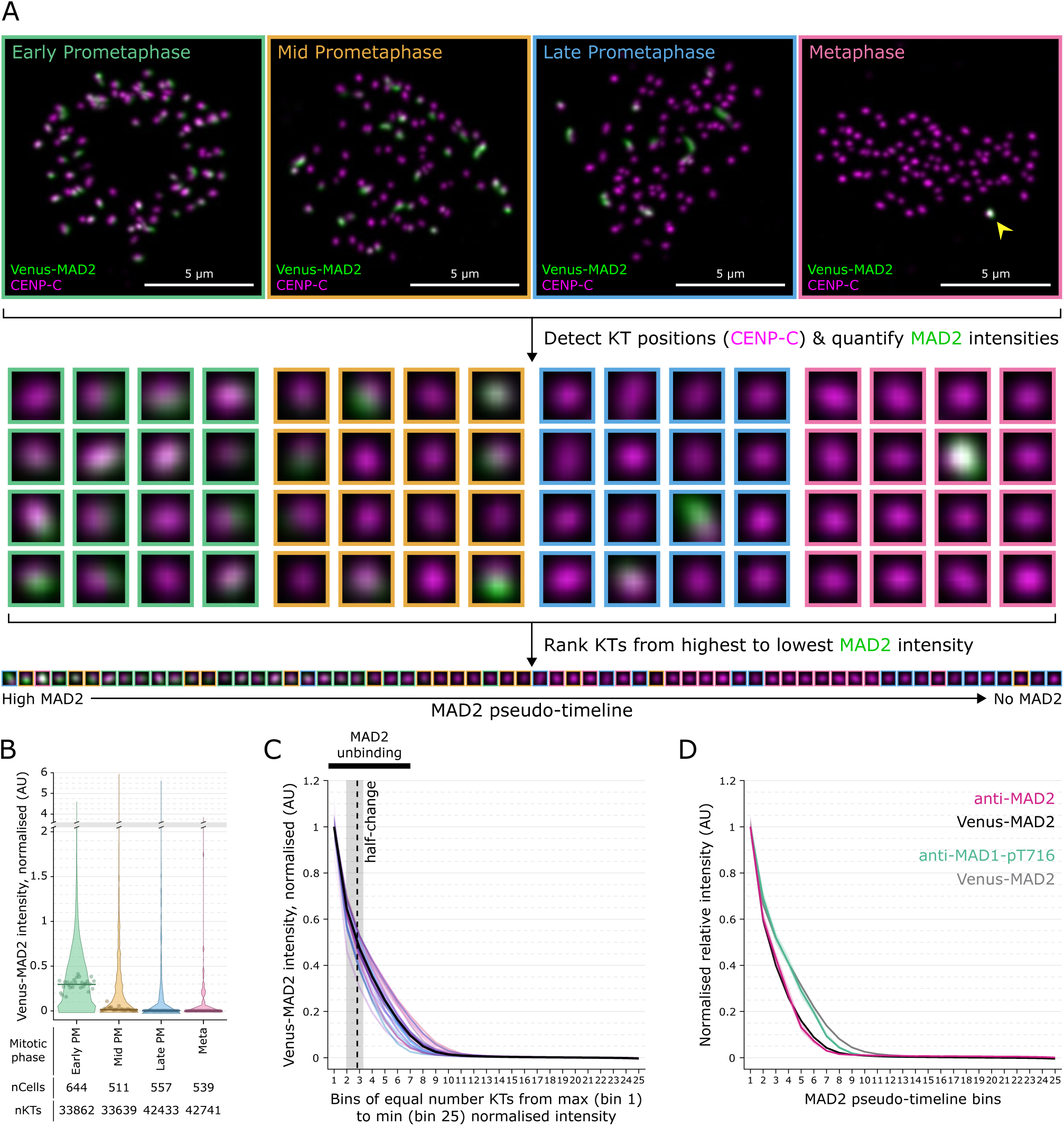
**Establishing a MAD2 pseudo-timeline to investigate SAC silencing.** All intensity measurements normalised to CENP-C. **(A)** Analysis pipeline. Top panel – Fixed unsynchronised RPE1 cells in early prometaphase (green box), mid prometaphase (yellow box), late prometaphase (blue box), and metaphase (pink box) were imaged. Endogenous Venus-MAD2 (green) and anti-CENP-C (magenta) are visualised. A high MAD2 kinetochore in the metaphase image is highlighted by a yellow arrowhead. Middle panel – A subset of kinetochores detected via CENP-C from each cell are visualised. Surrounding box colour indicates cell phase as in top panel. Lower panel – The same subset of kinetochores are ordered from maximum (left) to minimum (right) Venus-MAD2/CENP-C intensity. Surrounding box colour indicates cell phase as in top panel. **(B)** Violin plot of Venus-MAD2 intensity per mitotic phase. Median of each group is solid line, median of each repeat is represented by dot. Grey box represents y-axis break. **(C)** Pseudo-timelines created in individual experiments by endogenous Venus-MAD2; each colour represents a single experiment. nExpts = 27, nCells = 2247, nKTs = 152675. Black curve is full pseudo-timeline created from the combination of all data. Solid line represents the median intensity in each bin, shaded areas represent 95% confidence intervals of the median for each bin. Dotted vertical black line represents 50% change of Venus-MAD2 intensity of full pseudo-timeline (half-change), grey box represents range of Venus-MAD2 half-change across experiments. Solid black bar above graph represents 0-90% change of Venus-MAD2 intensity in full pseudo-timeline. Pseudo-timeline bins are represented by the x-axis, intensities are represented on the y-axis. **(D)**Anti-MAD2 antibody (pink; median and 95% confidence interval of median; nKTs = 13100, nCells = 164 from 2 repeats) and anti-MAD1-pT716 antibody (green; median and 95% confidence interval of median; nKTs = 11650, nCells = 166 from 2 repeats) intensities plotted on the MAD2 pseudo-timeline. Black and grey lines are the Venus-MAD2 intensities for anti-MAD2 antibody and anti-MAD1-pT716 antibody datasets, respectively.

### Early SAC silencing events can be ordered in MAD2 pseudo-time

The unbinding of MAD2 from kinetochores is thought to involve dephosphorylation of key kinetochore substrates, including KNL1 and BUB1 (Espert et al., 2014; London et al., 2012; Nijenhuis et al., 2014; Smith et al., 2019; Zhang et al., 2017), and Dynein-dependent transport along the spindle (Famulski et al., 2011; Gassmann et al., 2010; Howell et al., 2001; Ide et al., 2023; Silva et al., 2014). To determine whether these events occur at distinct MAD2 pseudo-times, we first analysed the kinetochore binding of BUBR1, which forms complexes with BUB1 and BUB3 on an array of phosphorylated MELT motifs in KNL1 (Krenn et al., 2014; Overlack et al., 2015; Yamagishi et al., 2012; Zhang et al., 2013; Zhang et al., 2015; Zhang et al., 2016). We found that unbinding of an endogenously labelled mEmerald-BUBR1 (Supplementary Figure 1) from kinetochores initiated at a similar pseudo-time but was right-shifted relative to the unbinding of MAD2 (Figure 2A, “MADs-BUBs lag”). BUBR1 was not completely removed, with a baseline of ∼15% retained at kinetochores (Figure 2A,B). We next followed the mitotic

**Figure 2:**
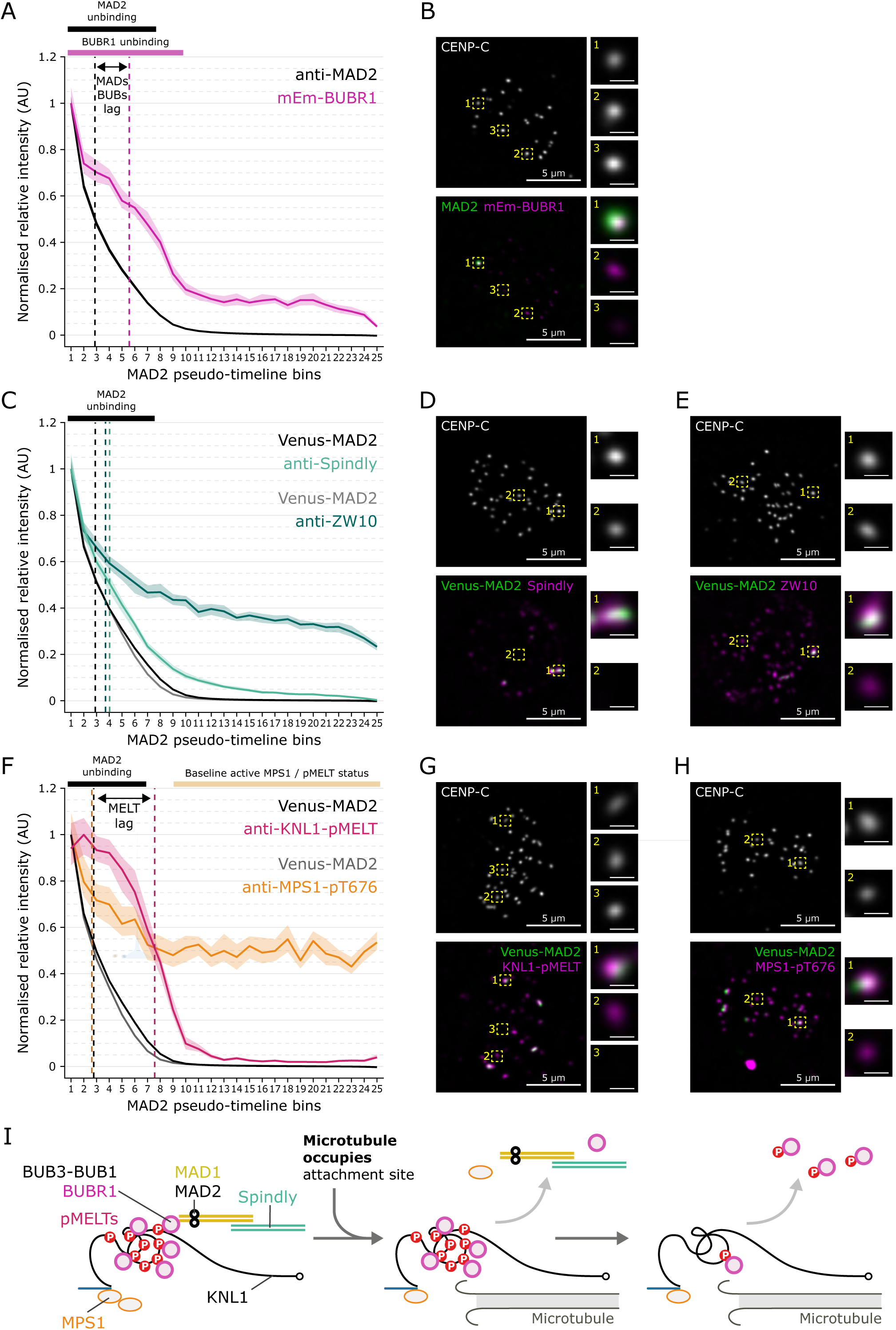
**Silencing initiates with unbinding of MAD2-Spindly and then disassembly of the KNL1 catalytic platform.** All intensity measurements normalised to CENP-C. **(A)** mEmerald-BUBR1 (pink; median and 95% confidence interval of median; nKTs = 14050, nCells = 183 from 2 repeats) intensities plotted on the MAD2 pseudo-timeline. Black line represents MAD2 intensity per bin for this dataset. Dotted vertical lines represent half-change of MAD2 (black) and mEmerald-BUBR1 (pink) intensities. Bars above graph represent represents 0-90% reduction in MAD2 intensity (black), and 0-90% reduction in mEmerald-BUBR1 intensity (pink). **(B, D, E, G, H)** Immunofluorescence of CENP-C and **(B)** MAD2 and mEmerald-BUBR1, **(D)** Venus-MAD2 and Spindly, **(E)** Venus-MAD2 and ZW10, **(G)** Venus-MAD2 and KNL1-pMELT 13/17, **(H)** Venus-MAD2 and MPS1-pT767. Partial projections of representative late prometaphase cells, with zoomed images of representative kinetochores. Maximum intensity projection of z-slices **(B)** 31-37, **(D)** 34-41, **(E)** 7-27, **(G)** 18-29, **(H)** 15-25. Zoomed image scale bars = 500 nm. Background centrosomic staining from anti-MPS1-pT676 antibody in **(H)** has been previously observed. **(C)** Spindly (light green; median and 95% confidence interval of median; nKTs = 12000, nCells = 168 from 2 repeats) and ZW10 (dark green; median and 95% confidence interval of median; nKTs = 12000, nCells = 157 from 2 repeats) intensities plotted on the MAD2 pseudo-timeline. Black and grey lines are Venus-MAD2 intensities for Spindly and ZW10 datasets, respectively. Dotted vertical lines represent half-change of Venus-MAD2 (black), ZW10 (dark green), and Spindly (green) intensities. Black bar above graph represents 0-90% reduction in MAD2 intensity. **(F)** KNL1-pMELT 13/17 (pink; median and 95% confidence interval of median; nKTs = 10900, nCells = 155 from 2 repeats) and MPS1-pT676 (orange; median and 95% confidence interval of median; nKTs = 12500, nCells = 179 from 2 repeats) intensities plotted on the MAD2 pseudo-timeline. Black and grey lines are Venus-MAD2 intensities for KNL1-pMELT and MPS1-pT676 datasets, respectively. Dotted vertical lines represent half-change of Venus-MAD2 (black), MPS1-pT676 (orange) and KNL1-pMELT (pink) intensities. Bars above graph represent 0-90% reduction in MAD2 intensity (black), and bins of minimum KNL1-pMELT and MPS1-pT676 intensities (light orange). **(I)** Schematic of early SAC silencing events. MAD1/MAD2 and Spindly leave when microtubules occupy attachment site, with concurrent partial loss of MPS1 and subsequent large-scale loss of KNL1-pMELTs and BUBR1.

Dynein adaptor Spindly, which is required for MAD2 stripping from the fibrous corona (Gassmann et al., 2010; Ide et al., 2023). In contrast to BUBR1, Spindly unbinding closely correlated with MAD2 and went to completion (Figure 2C,D). To gauge whether the whole fibrous corona followed these dynamics, we assessed the RZZ component ZW10 (Kops and Gassmann, 2020). This exhibited a similar half-change position to Spindly, but, unlike Spindly, did not fully unbind from kinetochores (Figure 2C,E). Thus, the initial phase of MAD2 unbinding coincides with motorised transport, perhaps in response to the first end-on contacts between microtubules and kinetochores.

The observed unbinding behaviour of BUBR1 suggested that dephosphorylation of KNL1-MELTs would also lag relative to MAD2 unbinding. To test this, we used a phospho-specific antibody to KNL1-pMELTs 13/17 (pT943/pT1155; Vleugel et al., 2015). Indeed, MELT phosphorylation remained high even when the first 50% of MAD2 is lost from kinetochores before declining to zero at approximately the same MAD2 pseudo-time that BUBR1 reaches steady state (Figure 2F,G). This supports the idea that initial MAD2 loss is not driven by KNL1-MELT dephosphorylation and the subsequent disassembly of the BUB3-BUB1-MAD1-MAD2 catalytic platform. Instead, this dephosphorylation may operate to reinforce SAC silencing and/or provide a window where MCC production can restart if microtubule attachments are lost.

Previous work suggested dephosphorylation of MELT motifs is favoured by the displacement of the MPS1 kinase from kinetochores (Hiruma et al., 2015; Ji et al., 2015; Pachis et al., 2019). This is proposed to be the result of competition between microtubules and MPS1 for a binding site on the Calponin Homology (CH) domain of NDC80 (Hiruma et al., 2015; Ji et al., 2015). We investigated how the kinetochore levels of phosphorylated MPS1 (which has higher kinase activity than its non-phosphorylated counterpart; Jelluma et al., 2008; Kang et al., 2007) relate to MAD2 pseudo-time. We found the MPS1-pT676 signal only reduced by 50%, with its half-change occurring coincident to that of MAD2, before MELT dephosphorylation had initiated (Figure 2F,H). This is consistent with the idea that MPS1 displacement promotes silencing. However, the sustained partial retention (even on kinetochores with no MAD2 signal) indicates that i) there is an active pool of MPS1 retained on kinetochores that cannot phosphorylate KNL1-MELTs, and ii) either there are insufficient microtubule contacts to fully displace MPS1 from NDC80, or there is a pool of MPS1 that is refractory to microtubule binding. The latter idea is supported by experiments in budding yeast where MPS1 does not completely leave even when the single microtubule is end-on attached to the kinetochore (Aravamudhan et al., 2015). From these initial experiments we can sketch a view of how early events are ordered during SAC silencing (in MAD2 pseudo-time; Figure 2I): the majority of MAD2 leaves first with Spindly, coincident with partial active MPS1 loss, followed by MELT dephosphorylation and dismantling of the BUB3-BUB1-BUBR1 catalytic platform.

### Eviction of MAD2 from kinetochores before substantial microtubule binding

We assume that MAD2 loss is instigated by initial microtubule contacts with the kinetochore. However, quantifying the number of microtubules needed for this and how microtubule occupancy levels affect MAD2 loss is challenging due to the number and density of microtubules in the spindle. We previously established that the supramolecular organisation of the NDC80 complexes at the kinetochore provides an indirect readout of microtubule binding: in unattached kinetochores (MAD2 positive) the ensemble of NDC80 complexes are highly disordered and jack-knifed, whilst in attached states (MAD2 negative) they become straight and ordered (Roscioli et al., 2020). We used this to determine when during the MAD2 pseudo-time NDC80 complexes change into the “attached” state. To do this, we calculated the Delta (inflation-corrected) distance between fluorescent spots formed by antibodies recognising CENP-C and the amino-terminus of NDC80 (NDC80-N; DeLuca et al., 2006). At early MAD2 pseudo-time, the Delta between CENP-C and NDC80-N was 15 nm and increased to 75 nm at later MAD2 pseudo-time. Notably, we discovered that the transition was switch-like and only initiated once 80% of MAD2 had unbound (Figure 3A,B). This suggests that the NDC80 ensemble straightens in a coordinated manner. Moreover, by measuring the distance between sister kinetochores, we found that the inter-sister tension generation closely followed the NDC80 switch (Figure 3C,D). Thus, high microtubule occupancy and tension (biorientation) are co-incident in MAD2 pseudo-time. The substantial lag between the half-change points of MAD2 unbinding and the NDC80 switch suggests that initiation of MAD2 loss must be in response to low microtubule occupancy of the kinetochore. This finding is supported by experiments showing that ∼50% microtubule occupancy is sufficient to silence the SAC (Dudka et al., 2018) and that ∼20% occupancy correlates with loss of MAD1 in living cells (Kuhn and Dumont, 2019). We speculate that perhaps only a single microtubule is necessary to initiate SAC silencing from an ensemble kinetochore. The mechanism underlying the necessary cooperativity will be important to investigate.

**Figure 3:**
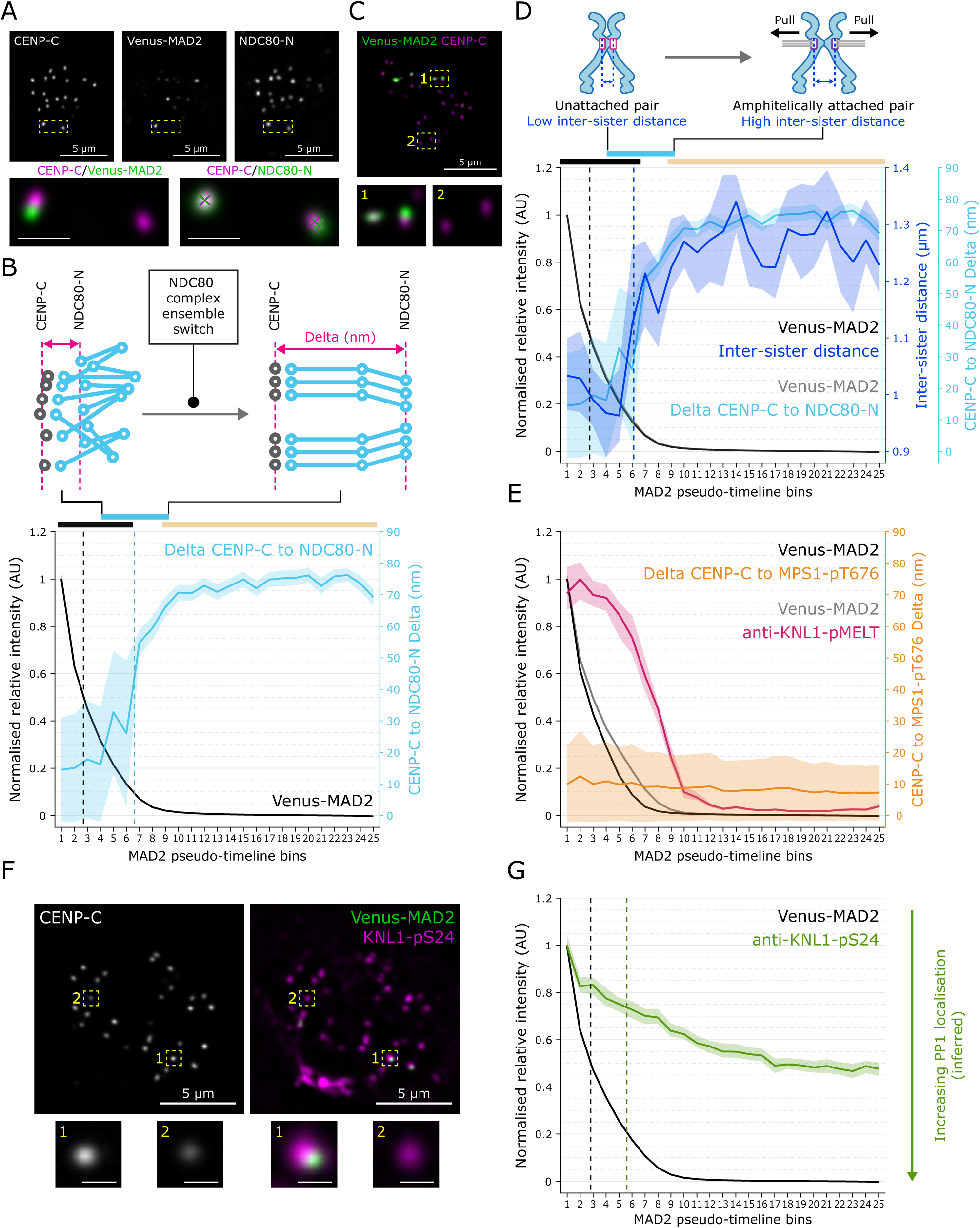
**Reconfiguration of the kinetochore outer plate happens after substantial MAD2 unbinding.** All intensity measurements normalised to CENP-C. **(A)** Immunofluorescence of CENP-C, Venus-MAD2 and NDC80-N. Partial projection of representative late prometaphase cell, with zoomed images of pair of kinetochores with high (left) and low (right) MAD2, and representative low (left) and high (right) CENP-C to NDC80-N intra-kinetochore distance. Maximum intensity projection of z-slices 45-55. Zoomed image scale bars = 1 µm. Magenta and green crosses on right zoomed image represent centre of CENP-C and NDC80-N (9G3) spots, respectively. **(B)** ntra-kinetochore Delta between CENP-C and NDC80-N (light blue; mean and 95% confidence interval of mean; nKTs = 16925, nCells = 348 from 4 repeats) plotted on the MAD2 pseudo-timeline; black line represents Venus-MAD2 intensity per bin for this dataset. Dotted vertical lines represent half-change of Venus-MAD2 intensity (black) and CENP-C to NDC80-N Delta (light blue). Bars above graph represent 0-90% reduction in MAD2 intensity (black), 0-90% change in CENP-C to NDC80-N Delta (blue), and bins of minimum KNL1-pMELT and MPS1-pT676 intensities (light orange). Schematics above graph represent changes in NDC80 conformation and organisation that contribute to Delta increase. **(C)** Immunofluorescence of CENP-C and Venus-MAD2. Partial projection (z-slices 47-52) of late prometaphase cell with zoomed images of high Venus-MAD2 and low Venus-MAD2 pair. Zoomed image scale bars = 1 µm. **(D)** Inter-sister distance (dark blue; median and 95% confidence interval of median; nPairs = 3063, nCells = 653 from 8 repeats) compared to CENP-C to NDC80-N intra-kinetochore distance (light blue, from Figure 3B) plotted on the MAD2 pseudo-timeline; black and grey lines are the Venus-MAD2 intensities for inter-sister distance and CENP-C to NDC80-N distance datasets, respectively. Dotted vertical lines represent half-change of Venus-MAD2 intensity (black) and inter-sister distance (dark blue). Bars above graph represent 0-90% reduction in MAD2 intensity (black), 0-90% change in CENP-C to NDC80-N Delta (blue), and bins of minimum KNL1-pMELT and MPS1-pT676 intensities (light orange). Schematics above graph represent changes in kinetochore attachment that may give rise to increased inter-sister distance. **(E)** Intra-kinetochore Delta between CENP-C and MPS1-pT676 (gold, mean and 95% confidence interval; nKTs = 9275, nCells = 179 from 2 repeats) compared to KNL1-pMELT intensity (pink, from Figure 2F) relative to the MAD2 pseudo-timeline; black and grey lines are the Venus-MAD2 intensities for CENP-C to MPS1-pT676 Delta and KNL1-pMELT intensity datasets, respectively. **(F)** Immunofluorescence of CENP-C, Venus-MAD2, and KNL1-pS24. Partial projection of representative late prometaphase cell with zoomed images of representative kinetochores. Maximum intensity projection of z-slices 32-42. Zoomed image scale bars = 500 nm. **(G)** KNL1-pS24 (green; median and 95% confidence interval of median; nKTs = 19450, nCells = 254 from 3 repeats) intensities plotted on the MAD2 pseudo-timeline. Black line represents Venus-MAD2 intensity per bin for this dataset. Dotted vertical lines represent half-change of Venus-MAD2 (black) and KNL1-pS24 (green) intensities.

### A second pool of MPS1 is refractory to microtubule binding and SAC silencing

The lag between partial MPS1 unbinding and the NDC80 switch hints that these events are uncoupled. Nevertheless, the MPS1 population that is retained on the kinetochore could be bound to the amino terminal CH domain in NDC80 (Hiruma et al., 2015; Ji et al., 2015), and would therefore be expected to change position as microtubules bind and NDC80 complexes increase their order. Such spatial changes are implicated in SAC silencing at budding yeast kinetochores (Aravamudhan et al., 2015). However, the Delta between MPS1-pT676 and CENP-C was ∼10 nm and did not change along the MAD2 pseudo-timeline (Figure 3E). Thus, the persistent MPS1 population is not bound on the NDC80 CH domain at the same time as microtubules. Instead, we infer that there is a second pool of MPS1 located close to the CCAN-KMN junction (Polley et al., 2024; Yatskevich et al., 2024).

The persistent MPS1 pool cannot phosphorylate KNL1-MELTs. One possibility is that phosphatases are beginning to dominate at this MAD2 pseudo-time. As BUBR1 (and thus its associated PP2A-B56; Kruse et al., 2013) is already leaving the kinetochore, we hypothesised that PP1 may be counteracting the remaining MPS1. To investigate PP1, we assessed phosphorylation levels of Serine 24 (S24) in KNL1. Dephosphorylation of this Aurora B site allows PP1 recruitment to the N-terminus of KNL1 (Bajaj et al., 2018; Welburn et al., 2010); thus, the inverse of KNL1-S24 phosphorylation was used as a proxy for PP1 recruitment. We found that KNL1-S24 underwent gradual dephosphorylation, suggesting that rising PP1 recruitment and phosphatase activity may limit MELT phosphorylation (Figure 3F,G). We note that tension onset did not accelerate KNL1-S24 dephosphorylation (unlike NDC80-S55 – see Figure 4) suggesting that initial microtubule contacts are sufficient to initiate some kinetochore dephosphorylation.

**Figure 4:**
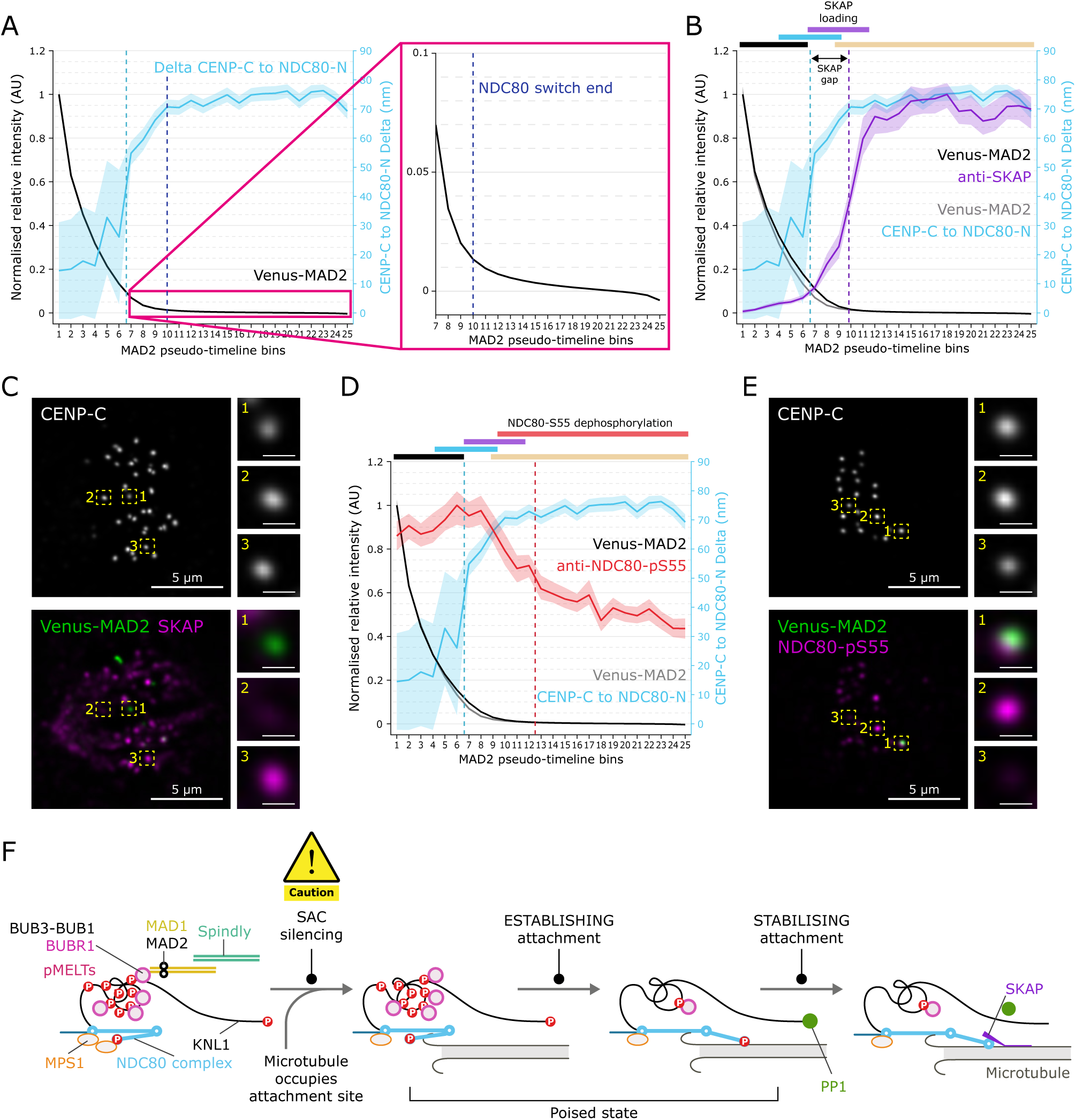
**NDC80 switch precedes recruitment of Astrin-SKAP and gradual attachment stabilisation through NDC80 tail dephosphorylation.** All intensity measurements normalised to CENP-C. **(A)** Data as Figure 3B, with zoomed segment of Venus-MAD2 intensity (pink box) after half-change of CENP-C to NDC80-N Delta (dotted vertical light blue line, full graph). Full change of CENP-C to NDC80-N Delta is marked on full graph and zoomed portion by dotted vertical dark blue line. **(B)** SKAP (purple; median and 95% confidence interval of median; nKTs = 11375, nCells = 167 from 2 repeats) intensity compared to CENP-C to NDC80-N Delta (light blue, from Figure 3B) plotted on the MAD2 pseudo-timeline; black and grey lines are the Venus-MAD2 intensities for SKAP intensity and CENP-C to NDC80-N Delta datasets, respectively. Dotted vertical lines represent half-change of CENP-C to NDC80-N Delta (blue) and SKAP intensity (purple). Bars above graph represent 0-90% reduction in MAD2 intensity (black), 0-90% change in CENP-C to NDC80-N Delta (blue), 10-90% change in SKAP intensity (purple), and bins of minimum KNL1-pMELT and MPS1-pT676 intensities (light orange). **(C, E)** Immunofluorescence of CENP-C, Venus-MAD2 and **(C)** SKAP, **(E)** NDC80-pS55. Partial projections of representative late prometaphase cells, with zoomed images of representative kinetochores. Maximum intensity projection of z-slices **(C)** 19-29, **(E)** 12-22. Zoomed image scale bars = 500 nm. **(D)** NDC80-pS55 (red; median and 95% confidence interval of median; nKTs = 15350, nCells = 238 from 3 repeats) intensity compared to CENP-C to NDC80-N Delta (light blue, from Figure 3B) plotted on the MAD2 pseudo-timeline; black and grey lines are the Venus-MAD2 intensities for NDC80-pS55 intensity and CENP-C to NDC80-N Delta datasets, respectively. Dotted vertical lines represent half-change of CENP-C to NDC80-N Delta (blue) and NDC80-pS55 intensity (red). Bars above graph represent 0-90% reduction in MAD2 intensity (black), 0-90% change in CENP-C to NDC80-N Delta (blue), 10-90% change in SKAP intensity (purple), full change of NDC80-pS55 intensity (red), and bins of minimum KNL1-pMELT and MPS1-pT676 intensities (light orange). **(F)** Schematic of attachment progression. SAC silences when microtubules occupy attachment site. Attachment is then established with NDC80 complex straightening, BUBs/pMELT loss and PP1 loading. The attachment is then stabilised by dephosphorylation of the NDC80 N-terminal tail and loading of the Astrin-SKAP complex.

### Stabilisation of microtubule-kinetochore attachments post-NDC80 switch

Following the NDC80 switch, the levels of MAD2 continue to decline (Figure 4A). Does this relate to continuing adaption of the kinetochore’s functional state, or is this simply unbinding of the final 2% of MAD2 from kinetochores? To investigate, we first assessed levels of the Astrin-SKAP complex component SKAP, which is known to mark mature end-on attachments (Dunsch et al., 2011; Fang et al., 2009; Friese et al., 2016; Kern et al., 2017; Mack and Compton, 2001; Manning et al., 2010; Schmidt et al., 2010). We found a clear switch-like recruitment of SKAP to kinetochores, which is right-shifted relative to the NDC80 switch (Figure 4B,C, “SKAP gap”). Kinetochores thus undergo further transitions between different functional states as the last fraction of MAD2 is lost. To further assess how attachments stabilise during this phase, we followed the dephosphorylation of Serine 55 (S55) located in the NDC80 N-terminal tail, a substrate for Aurora kinases (Ciferri et al., 2008; DeLuca et al., 2006; DeLuca et al., 2011; DeLuca et al., 2018). It is well established that phosphorylation of the NDC80 tail decreases the affinity of kinetochores for microtubules and operates to destabilise incorrect attachments during error correction (Alushin et al., 2012; Cheeseman et al., 2006; Ciferri et al., 2008; DeLuca et al., 2011; Guimaraes et al., 2008; Sarangapani et al., 2013; Welburn et al., 2010; Zaytsev et al., 2014; Zaytsev et al., 2015). We could detect no dephosphorylation during initial MADs/BUBs loss phase but upon inter-sister tension onset we observed continual dephosphorylation for the remainder of MAD2 pseudo-time (Figure 4D,E). This demonstrates that unbinding of MADs/BUBs, MELT dephosphorylation, and the NDC80 switch do not require dephosphorylation of NDC80-S55. Loss of the final 2% of MAD2 therefore provides insight into concurrent molecular changes at single kinetochores during this phase (which we term attachment maturation).

### Tension is dispensable for the NDC80 switch and SAC silencing

The MAD2 pseudo-timeline demonstrates how human kinetochores continue to adapt as the SAC is silenced, the key phases of which are summarised in Figure 4F. Our data implies that individual kinetochores commit to silencing their SAC signalling on initial contact with microtubules, with full attachment, stabilisation and maturation downstream events. This looks to be a hazardous approach as some kinetochores may well be improperly attached, requiring error correction to succeed before all other kinetochores silence their SAC signalling. It is therefore important to understand the relationship between error correction and SAC status at individual kinetochores. To investigate this, we treated cells with the Eg5 inhibitor monastrol to prevent centrosome separation, forming a monopolar spindle and making biorientation impossible (Kapoor et al., 2000). Around such monopoles sister kinetochores are enriched in syntelic and monotelic attachments which will undergo futile cycles of error correction, producing unattached kinetochores (Figure 5A). We could then construct a MAD2 pseudo-timeline for such kinetochores. We found that the maximum MAD2 levels were ∼3 times higher than in unperturbed (DMSO treated) cells (Figure 5B,C,E), presumably as kinetochores spend more time unattached. As expected, these kinetochores could not establish tension, with their inter-sister distance remaining close to rest length across MAD2 pseudo-time (Figure 5B).

**Figure 5:**
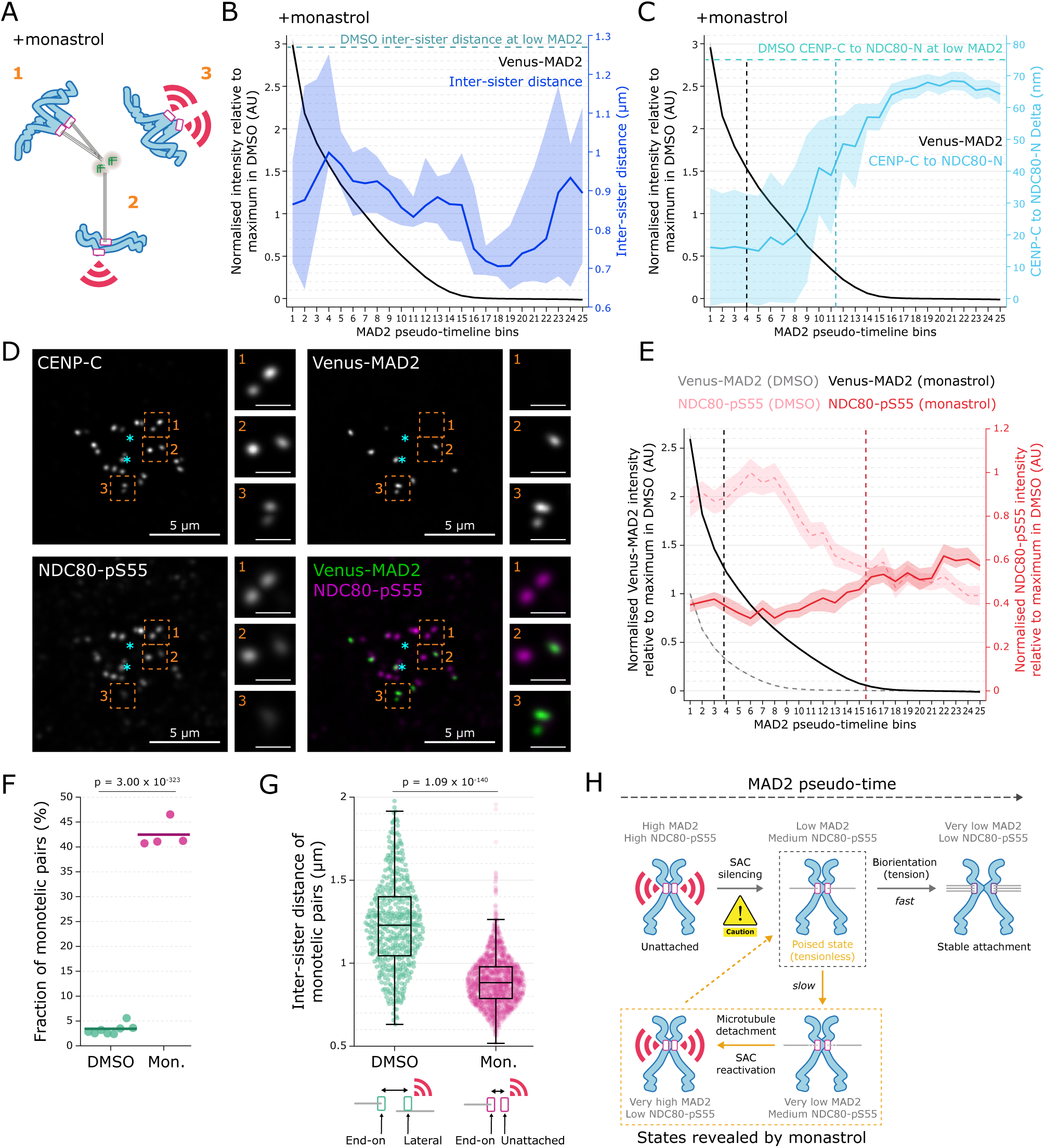
**Prevention of biorientation reveals hidden kinetochore states involved in error correction.** All intensity measurements normalised to CENP-C. **(A)** Schematic of attachment types in a monopolar system. 1. Syntelic attachment, 2. Monotelic attachment, 3. Unattached pair. **(B)** Inter-sister distance (dark blue; median and 95% confidence interval of median; nPairs = 599, nCells = 233 from 4 repeats) plotted on the monopolar spindle MAD2 pseudo-timeline; black line represents Venus-MAD2 intensities for this dataset. Dotted horizontal teal line represents average inter-sister distance of low MAD2 pairs in the bipolar spindle MAD2 pseudo-timeline. **(C)** Monastrol CENP-C to NDC80-N Delta (light blue; mean and 95% confidence interval of the mean; nKTs = 11175, nCells = 241 from 3 repeats) plotted on the MAD2 pseudo-timeline; black line represents Venus-MAD2 intensities for this dataset. Dotted horizontal turquoise line represents average CENP-C to NDC80-N Delta of low MAD2 kinetochores in the bipolar spindle MAD2 pseudo-timeline. Dotted vertical black lines represents half-change of Venus-MAD2 intensity (black) and CENP-C to NDC80-N Delta (blue) in monastrol treatment. **(D)** Immunofluorescence of CENP-C, Venus-MAD2, and NDC80-pS55 in a monastrol treated cell. Partial projection of a representative monastrol-treated cell (z = 38-52) with zoomed images of representative pairs in different attachment states. Pair 1 is syntelically attached, pair 2 is monotelically attached, pair 3 is unattached. Zoomed image scale bars = 1 µm. Cyan asterisks represent centrosome locations (z = 27 and 29). **(E)** NDC80-pS55 intensity in monastrol-treated cells (red; median and 95% confidence interval of median; nKTs = 14650, nCells = 259 from 3 repeats) compared to NDC80-pS55 intensity in DMSO-treated cells (dashed light pink, from Figure 4D) plotted on the MAD2 pseudo-timeline; black and dashed grey lines are the Venus-MAD2 intensities for monastrol and DMSO treatment datasets, respectively. Dotted vertical lines represent half-change of Venus-MAD2 (black) and NDC80-pS55 (red) intensities in monastrol treatment. **(F)** Percentage of monotelically attached pairs in DMSO and monastrol-treated cells. Dots represent individual repeats, solid lines represent overall percentages. DMSO: nPairs = 13665, nCells = 700. Monastrol: nPairs = 5206, nCells = 325. Fisher’s exact test was used for p-value. **(G)** Swarm plot of inter-sister distances of monotelically attached pairs in DMSO- and monastrol-treated cells. DMSO: nPairs = 471, nCells = 700; Monastrol: nPairs = 2212, nCells = 325. Each point is a pair. Wilcoxon rank sum test was used for p-value. Schematics show potential configurations that could result in the observed data. **(H)** Model of error correction. Kinetochores begin in an unattached state (top left). Microtubules occupy attachment sites, silencing the SAC. Kinetochores then enter a tensionless “poised” state (top centre), which can resolve in two ways: if the pair is bioriented, end-on attachments result in inter-sister tension, reducing Aurora B-mediated phosphorylation and permitting stable attachment (top right). Otherwise, if the tensionless state persists, Aurora B phosphorylates its substrates more extensively, destabilising attachments (bottom centre). Detachment results in reactivation of the SAC (bottom left) and permits another attempt at attachment.

Nevertheless, the MAD2 pseudo-timeline was similar to that seen in unperturbed cells and the NDC80 switch remains operational, albeit to a shorter final Delta than in unperturbed cells (Figure 5C), which we attribute to a lack of inter-sister tension. These data demonstrate that end-on attachment is sufficient for the NDC80 switch (and SAC silencing; Etemad et al., 2015; Kuhn and Dumont, 2017; Tauchman et al., 2015), with no requirement for biorientation and tension.

### Kinetochores silence the SAC before initiating error correction

Because biorientation is impossible in monastrol, and tension low, we expected to see elevated NDC80-pS55 signal across the MAD2 pseudo-timeline from increased Aurora B activity (McVey et al., 2021). However, we found that the NDC80-pS55 signal was inverted when compared to an unperturbed mitosis, with NDC80-S55 phosphorylation increasing even as the SAC silences (Figure 5D,E). We suspected that we are revealing error correction states of the kinetochore that are otherwise hidden in the total kinetochore population in an unperturbed mitosis. We confirmed this inverse correlation with visual inspection of sister kinetochore pairs and assigning attachment states based on MAD2 signal: where both sisters are MAD2 negative we infer syntely (Pair 1 in Figure 5A,D), while monotelic sisters had asymmetric MAD2 localisation (Pair 2) and unattached pairs had two MAD2 positive sisters (Pair 3). The signal in the NDC80-pS55 channel was clearly anti-correlated with MAD2 (Figure 5D), a result consistent with previous observations (Sobajima et al., 2023). This directly shows that the kinetochore can silence the SAC before initiating error correction. This is a risky strategy as the kinetochore relies on the success of error correction to prevent mis-segregation. Moreover, the switching of NDC80 into microtubule bound state is clearly not sufficient to allow its dephosphorylation and stabilisation of attachments – this step must also require a tension input.

### Error correction is rare in unperturbed mitosis

One prediction is that the monotelic attachment state is rarer in an unperturbed mitosis. To test this, we used the MAD2 signature identified in our monastrol experiments and classified monotelic pairs as those which had one sister with MAD2 intensity greater than 0.4 AU (before load-bearing attachments initiate on the control pseudo-timeline) and the other with MAD2 intensity less than 0.01 AU (where kinetochores have mature end-on attachments, marked by SKAP in the control pseudo-timeline). We found that monotelic pairs were 12-fold less frequent in a normal mitosis than monastrol (3.4% versus 42.5%; Figure 5F). These monotelic pairs, unlike those found in monastrol, were under tension (Figure 5G). This configuration can be explained by previous light and electron microscopy experiments, where monotelic pairs in a bipolar spindle were under tension with one sister end-on and the other laterally attached (Kapoor et al., 2006; Kuhn and Dumont, 2017). By contrast, the monotelic pairs in monastrol (one end-on attached, one unattached) would presumably arise from the completion of error correction.

### Integration of SAC silencing and error correction

Figure 5H proposes a model that reconciles our findings: Unattached kinetochores are positive for both MAD2 and NDC80-pS55 (top left). MAD2 unbinding proceeds rapidly following initial microtubule contacts. Live cell measurements suggest this happens on the order of several minutes in real time (Kuhn and Dumont, 2017; Sikirzhytski et al., 2018). This leads to a kinetochore which has lost MAD2 but is still positive for NDC80-pS55, active MPS1 and BUBs. We propose this to be a “poised state” of the kinetochore that allows for rapid reactivation of the SAC if there is a loss of attachment (top centre). We have previously proposed that such responsiveness is required for an optimal SAC (McAinsh and Kops, 2023). In the situation where tension can be generated, then the “poised” kinetochore would quickly move into the stable and mature attachment state (top right). When biorientation is not possible and tension is low, the phosphorylation of Aurora substrates will be favoured, resulting in error correction (bottom centre) and reactivation of the SAC. We suspect that the unattached kinetochores in monastrol are the product of error correction (bottom left). One puzzle is that these have very low NDC80-pS55 compared to unattached kinetochores in unperturbed cells, which have very high NDC80-pS55. It is possible that this reflects differences between kinetochores that have never bound microtubules before versus those that are the product of error correction cycle(s); indeed, it has previously been demonstrated that kinetochores which lose microtubule attachments are incapable of returning to a maximal phosphorylation state (DeLuca et al., 2011).

### Kinetochores are adaptive machines

Here we create “MAD2 pseudo-time” and exploit this to reveal how human kinetochores transition between multiple functional states during SAC silencing. Our data show that SAC silencing is not a simple ON/OFF switch with multiple co-incident molecular events – rather, the number of MAD2 molecules encodes and unveils information on the underlying functional state of the kinetochore. This approach allows individual molecular events to be ordered and reveals several new phases of kinetochore biology (see model in Figure 4F). The transitions between states also display unique dynamics, with evidence of both switch-like (e.g. NDC80 ensemble straightening or SKAP loading) and more gradual events (e.g. Aurora substrate dephosphorylation). This shifts our understanding of how molecular events are causally related, i.e. NDC80 complex binding to microtubules cannot trigger initial loss of MAD2. To what extent an event gates a subsequent event can now be investigated. Kinetochores thus operate as molecular machines that process inputs, then adapt and update their functional state to ensure accurate chromosome segregation.

## Materials and Methods

### Key Resources Table

| Reagent or Resource | Source | Identifier |
| --- | --- | --- |
| **Antibodies** |  |  |
| **Immunofluorescence** |  |  |
| Guinea Pig polyclonal anti-CENP-C (1:2000) | MBL International | Cat# PD030, RRID:AB_10693556 |
| Rabbit polyclonal anti-MAD2 (1:500) | Bethyl | Cat# A300-301A, RRID:AB_2281536 |
| Rabbit polyclonal anti-MAD1-pT716 (1:1000) | Allan et al., 2020 |  |
| Rabbit polyclonal anti-Spindly (1:200) | Bethyl | Cat# A301-354A, RRID:AB_937753 |
| Rabbit polyclonal anti-ZW10 (1:200) | Abcam | Cat# ab21582, RRID:AB_779030 |
| Rabbit polyclonal anti-KNL1-pT943/pT1155 [MELT 13/17] (1:2000) | Vleugel et al., 2015 |  |
| Rabbit polyclonal anti-MPS1-pT676 (1:4000) | Jelluma et al., 2008 |  |
| Mouse monoclonal anti-Hec1 [9G3] (1:4000) | Abcam | Cat# ab3613, RRID:AB_303949 |
| Rabbit polyclonal anti-KNL1-pS24 (1:2200) | Welburn et al., 2010 |  |
| Rabbit polyclonal anti-SKAP (1:400) | Atlas Antibodies | Cat# HPA042027, RRID:AB_10797378 |
| Rabbit polyclonal anti-phospho Hec1 Ser55 (1:500) | Thermo Fisher Scientific | Cat# PA5-34980, RRID:AB_2552328 |
| Goat anti-guinea pig AlexaFluor 488 (1:500) | Invitrogen | Cat# A-11073, RRID:AB_2534117 |
| Goat anti-guinea pig AlexaFluor 594 (1:500) | Invitrogen | Cat# A-11076, RRID:AB_2534120 |
| Goat anti-guinea pig AlexaFluor 647 (1:500) | Invitrogen | Cat# A-21450, RRID:AB_2535867 |
| Goat anti-rabbit AlexaFluor 647 (1:500) | Invitrogen | Cat# A-21245, RRID:AB_2535813 |
| Goat anti-mouse AlexaFluor 647 (1:500) | Invitrogen | Cat# A32728, RRID:AB_2633277 |
| **Immunoblotting** |  |  |
| Rabbit polyclonal anti-BUBR1 (1:500) | Bethyl | Cat# A300-386A, RRID:AB_386097 |
| Mouse monoclonal anti-CDC20 (1:500) | Santa Cruz Biotechnology | Cat# sc-13162, RRID:AB_628089 |
| Mouse monoclonal anti-Vinculin (1:1000) | Santa Cruz Biotechnology | Cat# sc-73614, RRID:AB_1131294 |
| IRDye800CW donkey anti-rabbit (1:10,000) | LI-COR Biosciences | Cat# 926-32213, RRID:AB_621848 |
| IRDye680RD donkey anti-mouse (1:10,000) | LI-COR Biosciences | Cat# 926-68072, RRID:AB_10953628 |
| **Chemicals** |  |  |
| DMSO | Sigma-Aldrich | Cat# D2438 |
| Monastrol | Sigma-Aldrich | Cat# M8515 |
| VECTASHIELD Antifade Mounting Medium with DAPI | Vector Laboratories | Cat# H-1200-10 |
| **Software and algorithms** |  |  |
| KiT 2.4.0 | This paper | https://github.com/cmcb-warwick/KiT |
| Bayesian Euclidean distance correction algorithm | Roscioli et al., 2020 | https://github.com/cmcb-warwick/KiT |
| MATLAB (2022a) | MathWorks |  |
| Huygens 4.2 (Deconvolution software) | SVI |  |
| Volocity v6.3.1 (UltraVIEW microscope software) | PerkinElmer |  |
| Fiji | Open source |  |

### Cell culture

Human hTERT immortalised non-transformed diploid retinal pigment epithelial (RPE1) cells (ATCC code), RPE1 Venus-MAD2 (*MAD2L1* Venus/+; Collin et al., 2013), RPE1 3xFLAG-mEmerald-BUBR1+/+ (this paper) were grown in DMEM/F-12 medium containing 10% Fetal Bovine Serum, 100 U/mL penicillin, 100 µg/mL streptomycin, and 2 mM L-glutamine. Cells were maintained at 37°C with 5% CO_2_ in a humidified incubator.

### Cell line creation

To tag BUBR1, 3 x 10^6^ RPE FRT/TR cells were transfected with 3 µg of modified PX466 ‘All-in-One’ plasmid containing Cas9D10A-T2A-mRuby and gRNAs targeting *BUB1B* (5’-CTACCAGGCCGTGTCCTCTC-3’ and 5’-GGCTCCGCTCGGAGCAGTCG-3’). The all-in-one plasmid was co-transfected alongside 3 µg of repair plasmid that contained 3xFLAG-mEmerald-GGGGS flanked by ∼500 bp arms that were homologous to the genomic region that surrounds the Cas9 cutting site. 72 hr post transfection, 50,000 mRuby-positive cells were sorted into a 1 cm well and expanded for one week. mEmerald-positive cells were then single-cell sorted into 96-well plates. The mEmerald tag was identified through PCR screening using primers 5’-TCAGTCAAGGCTAGTCCCCT-3’ and 5’-TGGCATAGCACCTGCCATTT-3’.

Further PCRs were conducted on the clones of interest using the following primer sets: 5’-TCAGTCAAGGCTAGTCCCCT-3’ and 5’-TCTAGGTATTGCGGGCACCA-3’, 5’-TCAGTCAAGGCTAGTCCCCT-3’ and 5’-GTGGCTGTTGTAGTTGTACTCC-3’, and 5’-CCGAAGGCTACGTCCAGG-3’ and 5’-TCTAGGTATTGCGGGCACCA-3’, and the products sequenced. As only heterozygous clones were identified, the transfection, sorting and screening process was repeated on two of these clones using a modified guide (5’-CTACCAGGCCGTGTTCTCTC-3’ paired with 5’-GGCTCCGCTCGGAGCAGTCG-3’) which targeted a SNP that was initially preventing the editing of one allele. All transfections were performed by electroporation using a Neon Transfection System (Invitrogen) with two pulses at 1400 V for 20 ms.

### Protein extraction

Adherent cells were trypsinized and incubated in lysis buffer (150 mM NaCl, 50 mM Tris pH 7.4, 0.5% NP-40) supplemented with HALT protease and phosphatase inhibitor cocktail (Thermo Scientific) for 30 minutes at 4°C. Lysates were then clarified by centrifugation for 20 minutes at 14,000 x g and 4°C, and quantified using Bradford Reagent (Bio-Rad Laboratories) according to the manufacturer’s instructions.

### Immunoblotting

Fifty micrograms of cell lysates were separated through SDS-PAGE on a NuPAGE 4-12% Bis-Tris gel (Invitrogen) and transferred to an Immobilon-FL polyvinylidene fluoride membrane (IPFL00010, Millipore). The membrane was blocked using 5% milk, 0.1% Tween in PBS, then incubated at 4°C overnight with primary antibodies in 2.5% milk, 0.1% Tween in PBS. The next day, the membrane was washed with 0.1% Tween in PBS, and incubated for 1 hour at room temperature with secondary antibodies in 2.5% milk, 0.1% Tween in PBS. A LI-COR Odyssey CLx Imager (LI-COR Biosciences) was used for visualisation.

### Drug treatments

Cells were plated on #1.5 22×22 mm glass coverslips 24-48h prior to drug treatment. Cells were treated with 100 µM monastrol diluted in DMSO for 4h or with 0.1% DMSO for 4h as a control.

### Indirect immunofluorescence

Drug-treated and control cells were fixed in PTEMF (20 mM PIPES buffer, pH 6.8; 0.2% Triton X-100; 10 mM EGTA buffer, pH 7.0; 1 mM MgCl_2_; 4% formaldehyde) for 10 minutes, washed with PBS 3 times for 5 minutes, then incubated in PBS containing 3% BSA to reduce non-specific antibody binding for 30 minutes. Cells were then incubated with primary antibodies for 1 hour, washed with PBS 3 times for 5 minutes, and incubated with secondary antibodies for 30 minutes. All antibodies were diluted in PBS containing 3% BSA. Cells were washed in PBS 3 times for 5 minutes, then mounted in DAPI-containing VectaShield mounting medium, sealed with nail polish, and stored at 4°C. All steps were performed at room temperature with samples shielded from light.

### Image collection

Image stacks were acquired using a confocal spinning disk microscope (VOX UltraVIEW, PerkinElmer) equipped with Nikon Plan Apochromat 100x 1.40 NA oil-immersion objective and SCMOs Hamamatsu Orca-Flash4.0 v3 camera (16-bit), controlled by Volocity 6.3.1 (PerkinElmer) running on a Windows 10 64-bit PC. For all images, an xy area of 512 x 512 pixels was collected, with 69.3 nm effective pixel size. Image stacks were acquired using the 488, 560, 640, and 405 nm lasers. For samples, image stacks were acquired over 61 z-slices separated by 200 nm; for channel alignment, image stacks were acquired over 121 z-slices separated by 100 nm. Laser power and exposure time were set in order that maximum kinetochore signals were around 300 units above background.

### Image analysis

Images were exported from Volocity in OME.TIFF format (Open Microscopy Environment). Microscope-specific point spread functions (PSFs) were calculated in Huygens 4.2 PSF distiller from 100 nm TetraSpeck fluorescent microspheres. These PSFs were used to deconvolve images in Huygens 4.2, which were exported in r3d format. Deconvolved images were read into MATLAB using the bfmatlab toolbox (Open Microscopy Environment). KiT (Kinetochore Tracking) v2.4.0 (running in MATLAB) was used for kinetochore detection, spot refinement, manual sister pairing (performed prior to 3D intra- and inter-kinetochore measurements only), spot quality control (manual), intensity measurements, and 3D intra- and inter-kinetochore distance measurements (Armond et al., 2016; Dudka et al., 2018). Initial alignment of channels was carried out using images taken from a reference slide (RPE1 cells probed with anti-CENP-C, stained with a mix of AlexaFluor 488, 594, and 647-labelled secondary antibodies) on the day of image acquisition. Kinetochore spots were first detected using CENP-C signal in the 561 nm channel, with spots in the secondary and tertiary channels subsequently detected where appropriate. If performing intra-kinetochore or inter-sister distance measurements, kinetochores were manually paired using CENP-C spots. CENP-C spots, and secondary/tertiary channel spots where appropriate, were manually filtered to remove overlapping or non-Gaussian spots. Secondary/tertiary marker spot positions were corrected to make the average intra-kinetochore distance per cell in x-, y-, and z-directions equal to 0 (Armond et al., 2016; Dudka et al., 2018). Intensities in each channel were measured in a 300 nm circular radius centred at the channel offset-corrected CENP-C spot location, background subtracted, and normalised using the background-subtracted CENP-C signal. Inflation-corrected intra-kinetochore Delta distances per bin (see below for binning details) were computed from 3D Euclidean intra-kinetochore distances per bin using BEDCA v1.0 running in MATLAB (Germanova et al., 2021; Roscioli et al., 2020). Statistical tests were performed in MATLAB.

### MAD2 pseudo-timeline graphs

In each experimental repeat, kinetochores were ordered from maximum to minimum relative (Venus-)MAD2 intensity. These kinetochores were then assigned to 25 equal-sized bins, each representing 4% of kinetochores. The median intensity value of each bin was calculated, and intensities were normalised so the maximum median bin intensity of MAD2 (and secondary marker where appropriate) was equal to 1 for control experiments; intensities from monastrol treatment experiments were normalised to their paired control experiment. Experimental repeats were combined post-normalisation, with kinetochores re-ordered and re-binned based on their normalised relative MAD2 intensities. For inter-sister distances, only sisters whose MAD2 intensities were in adjacent bins were accepted for plotting; the mean intensity rank of the pair was used for bin placement when plotting. Confidence intervals of median were calculated as described by Ialongo (2019).

## Acknowledgements

We thank Iain Cheeseman for anti-KNL1-pS24 antibody, Geert Kops for anti-KNL1-pMELT 13/17 and anti-MPS1-pT676 antibodies, and Adrian Saurin for anti-MAD1-pT716 antibody. We also thank Warwick’s Computational and Advanced Microscopy Development Unit (CAMDU) for microscopy support and Nina Pučeková for comments on the manuscript. C.C.C. and T.G. were funded by the Medical Research Council Doctoral Training Partnership in Interdisciplinary Biomedical Research (MR/N014294/1 and MR/J003964/1, respectively). A.D.M. and S.T.P. are supported by a Wellcome Discovery Award (216605/Z/22/Z). C.C. and J.P. are supported by a Wellcome Trust Investigator Award (209470/Z/17/Z).

## Author Contributions

C.C.C.: Conceptualisation, methodology, investigation, formal analysis, writing – reviewing and editing, figure preparation. T.G.: Conceptualisation (initiated project with A.D.M. and N.J.B.), methodology, investigation (including the initial experiments to establish the MAD2 pseudo-timeline). S.T.P.: Formal analysis, Reviewing. C.C.: Resources (generation of mEmerald-BUBR1 +/+ homozygous knock-in cells). J.P.: Supervision. N.J.B.: Conceptualisation, supervision. A.D.M.: Conceptualisation, supervision, formal analysis, Writing – original draft preparation, reviewing and editing, figure preparation, funding acquisition.

---

**Supplementary Figure 1:**
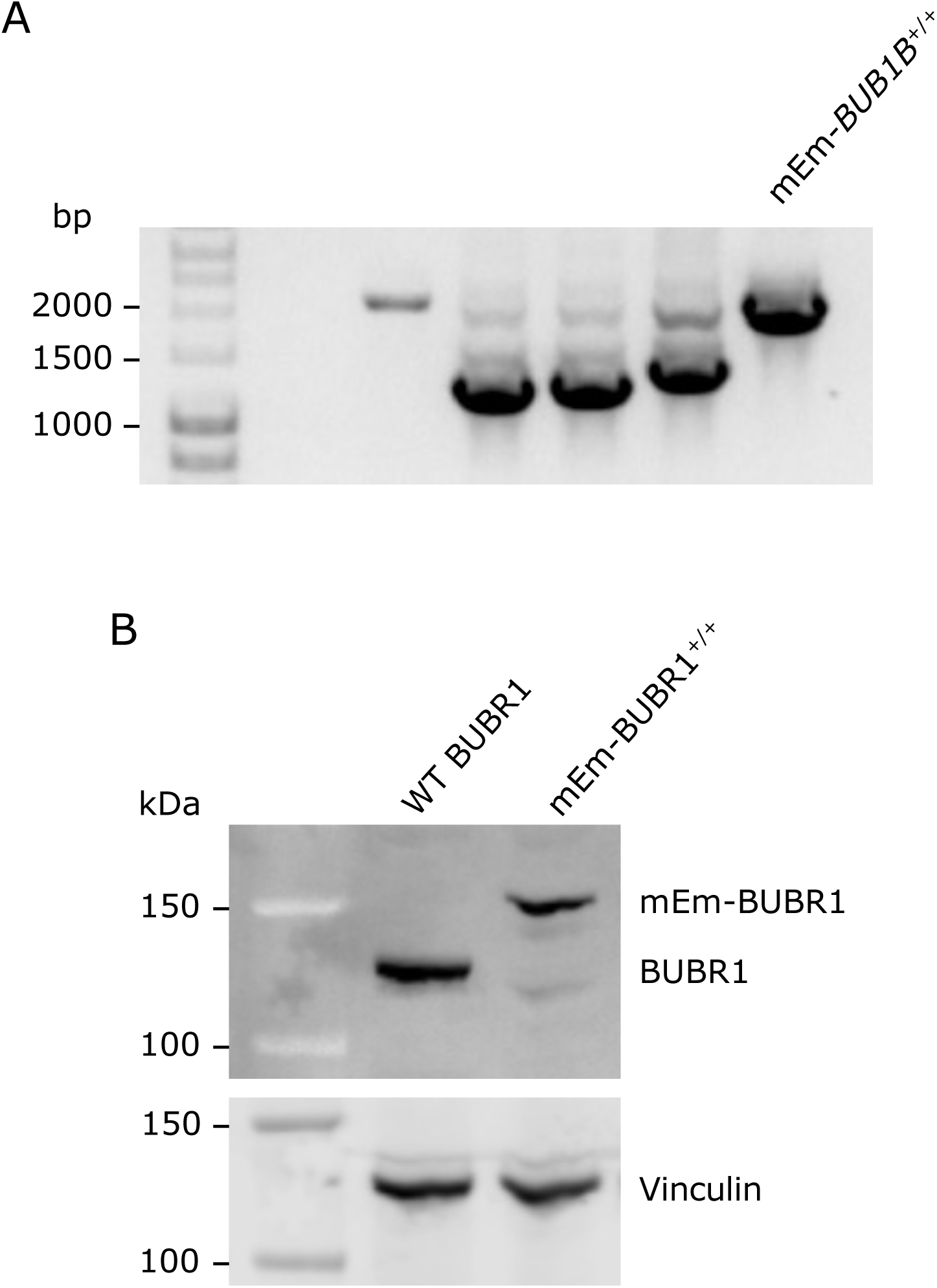
Validation of RPE1 mEmerald-BUBR1+/+ cell line. **(A)** Agarose gel showing a PCR screen of clones. A band at 1275 bp indicates the presence of wild-type *BUB1B*, a band at 2058 bp indicates that *BUB1B* has been edited to include 3x-FLAG-mEmerald-linker after the start codon. **(B)** Western blot using anti-BUBR1 and anti-Vinculin against lysates from parental RPE1 FRT/TR cells or from RPE1 FRT/TR mEmerald-BUBR1+/+ cells.

